# Fluidic microactuation of flexible electrodes for neural recording

**DOI:** 10.1101/155937

**Authors:** Flavia Vitale, Daniel G. Vercosa, Alexander V. Rodriguez, Sushma Sri Pamulapati, Frederik Seibt, Eric Lewis, J. Stephen Yan, Krishna Badhiwala, Mohammed Adnan, Gianni Royer-Carfagni, Michael Beierlein, Caleb Kemere, Matteo Pasquali, Jacob T. Robinson

## Abstract

Ultra-flexible microelectrodes that can bend and flex with the natural movement of the brain reduce the inflammatory response and improve the stability of long-term neural recordings.^1-5^ However, current methods to implant these highly flexible electrodes rely on temporary stiffening agents that increase the electrode size^6-10^ thus aggravating neural damage during implantation, which leads to cell loss and glial activation that persists even after the stiffening agents are removed or dissolve.^11-13^ A method to deliver thin, ultra-flexible electrodes deep into neural tissue without increasing the stiffness or size of the electrodes will enable minimally invasive electrical recordings from within the brain. Here we show that specially designed microfluidic devices can apply a tension force to ultra-flexible electrodes that prevents buckling without increasing the thickness or stiffness of the electrode during implantation. Additionally, these “fluidic microdrives” allow us to precisely actuate the electrode position with micron-scale accuracy. To demonstrate the efficacy of our fluidic microdrives, we used them to actuate highly flexible carbon nanotube fiber (CNTf) microelectrodes^11,14^ for electrophysiology. We used this approach in three proof-of-concept experiments. First, we recorded compound action potentials in a soft model organism, the small cnidarian *Hydra*. Second, we targeted electrodes precisely to the thalamic reticular nucleus in brain slices and recorded spontaneous and optogenetically-evoked extracellular action potentials. Finally, we inserted electrodes more than 4 mm deep into the brain of rats and detected spontaneous individual unit activity in both cortical and subcortical regions. Compared to syringe injection, fluidic microdrives do not penetrate the brain and prevent changes in intracranial pressure by diverting fluid away from the injection site during insertion and actuation. Overall, the fluidic microdrive technology provides a robust new method to implant and actuate ultra-flexible neural electrodes.

---

Chronically-implanting electrodes enables measurement of the spiking activity of individual neurons in freely-behaving animals, which illuminate the neural processes underlying learning, movement, perception and cognition, in healthy and diseased states. Over the past few years these chronically-implanted microelectrodes have led to breakthrough discoveries and technological innovations including control of robotic end effectors,^15,16^ restoration of the cortical control of the upper limbs,^17^ discovery of fundamental mechanisms underlying cognitive processes,^18,19^ and identification of unique neural firing patterns associated with epileptic activity.^20-22^

Despite their foundational role in systems neuroscience and brain-machine interfaces, existing microelectrode technologies have significant challenges. Electrodes have traditionally been manufactured using metal wires or micromachined silicon. The roughly 10^8^-fold stiffness mismatch between the electrode material and the soft host brain tissue causes acute and chronic injury that results in extensive neuronal death, formation of gliotic encapsulation and isolation of the recording sites from the neuronal bodies.^23,24^ Moreover, relative micromotion between the tissue and the implant can exacerbate the strain-induced inflammatory reaction and cause the recording site to drift from its original position.^25,26^ Thus, creating electrodes from highly flexible materials holds great promise for improving the electrode-brain interface.

Engineered flexible substrates and ultrasmall microwires approximating the diameter of individual cells have been shown to significantly mitigate the chronic injury and improve signal quality and longevity of neural recordings.^1-5,27^ Recently, we demonstrated that flexible carbon nanotube fiber (CNTf) microelectrodes are effective for neuromodulation and chronic recording applications, while eliciting minimal foreign body reaction when compared to metal implants.^11^ The major drawback of these novel compliant materials lies in insertion – when the unsupported length of the electrodes is longer than a critical limit (typically order 1-2 mm for state of the art electrodes)^28,29^ they buckle rather than enter the brain tissue. This challenge has been solved by temporarily increasing the stiffness of the electrode.^10^ For example, flexible electrodes can be stiffened through attachment to a rigid probe during insertion^4,6,11^ or by overcoating with a hydrogel sheath^7-9^ that dissolves minutes after implantation. However, the increased rigidity and footprint of the stiffener-electrode assembly can aggravate the acute and chronic injury, damaging or destroying nearby neurons and breaching the blood brain barrier.^11-13^ Recent work^5,30^ demonstrated that macroporous 3-D mesh electrodes with cellular-scale feature sizes (~20 μm) and mechanical flexibility 10^7^ times greater than conventional silicon greatly improve the integration of neural electrodes with brain tissue and are capable of intraoperative and chronic recordings *in vivo*. However, to implant these devices a rigid syringe with a diameter of 650 μm must be inserted into the cortex.^5,30^

A new technology to insert and precisely position flexible, cellular-scale electrodes without temporary stiffening would significantly improve the robustness and utility of flexible electrodes while minimizing excess damage to the brain from stiffeners. To reach this goal we developed a novel strategy to implant and microactuate flexible microelectrodes without using external supports or stiffening agents. Instead, we use viscous fluid flow in a microfluidic channel to maintain tension in the electrode structure – effectively stiffening it without increasing the implant footprint. Using microfluidic vent channels, we can then divert nearly all the fluid away from the point of electrode solution. The result is a fluidic microdrive technology which enables the accurately-controlled insertion of flexible electrodes. As a proof-of-concept, we demonstrate electrophysiological recordings from ultraflexible cellular-scale electrodes following fluidic implantation into small organisms, acute mouse brain slices, and the brain of anesthetized rats

## Results

### CNTf Electrodes

Cellular-scale CNTf microwires are promising neural electrodes for stimulation and recording^11^ whose flexibility presents a significant challenge for neural implantation. For this study, we used CNTf microwires with diameter between 12 and 25 μm insulated with either a conformal bilayer of 50 nm Al_2_O_3_ and 25 nm of HfO_2_^31^ or 1.2 to 3.3 μm of parylene C (Supplementary Figure S1). After the insulation step, we measured the bending stiffness of the parylene-coated CNTf microelectrodes to be 0.23 x 10^-9^ Nm^2^ for the 12 μm diameter probes (post-insulation diameter 18.3 ± 0.7 μm, mean ± s.d., n = 3) and 1.08 x 10^-9^ Nm^2^ for the 25 μm diameter probes (post-diameter thickness 28.9 ± 0.6 μm, n = 3), which is comparable to the flexibility of polyimide probes that have been shown to require stiffening supports for successful surgical implantation^32^ (Supplementary Figure S3). As the penetration of water and ions through the insulating coatings is one of the major causes leading to the failure of chronic neural implants and it is still an active area of investigation,^33,34^ we performed leakage current and impedance spectroscopy measurements before and after 90° bending the parylene-coated CNTf and found no evidence of acute breakdown of the insulating layer (Supplementary Figure S2). To expose the conductive CNTf core at the ends of the electrode, we cut the electrode using a razor blade or focused ion beam (FIB).

### Fluidic microdrive fabrication

We fabricated microfluidic microdrives from two layers of polydimethylsiloxane (PDMS) using conventional replica-molding techniques.^35^ As shown in Figure 1, fluid flows through the channel containing the electrode, but is diverted away from the implantation site to minimize tissue damage. We manually placed CNTf microelectrodes in the center of the channel before plasma bonding the channels to a glass substrate. After the PDMS device was bonded, plastic tubing was plugged into the flow ports that connect the device to the flow control system.

**Figure 1.**
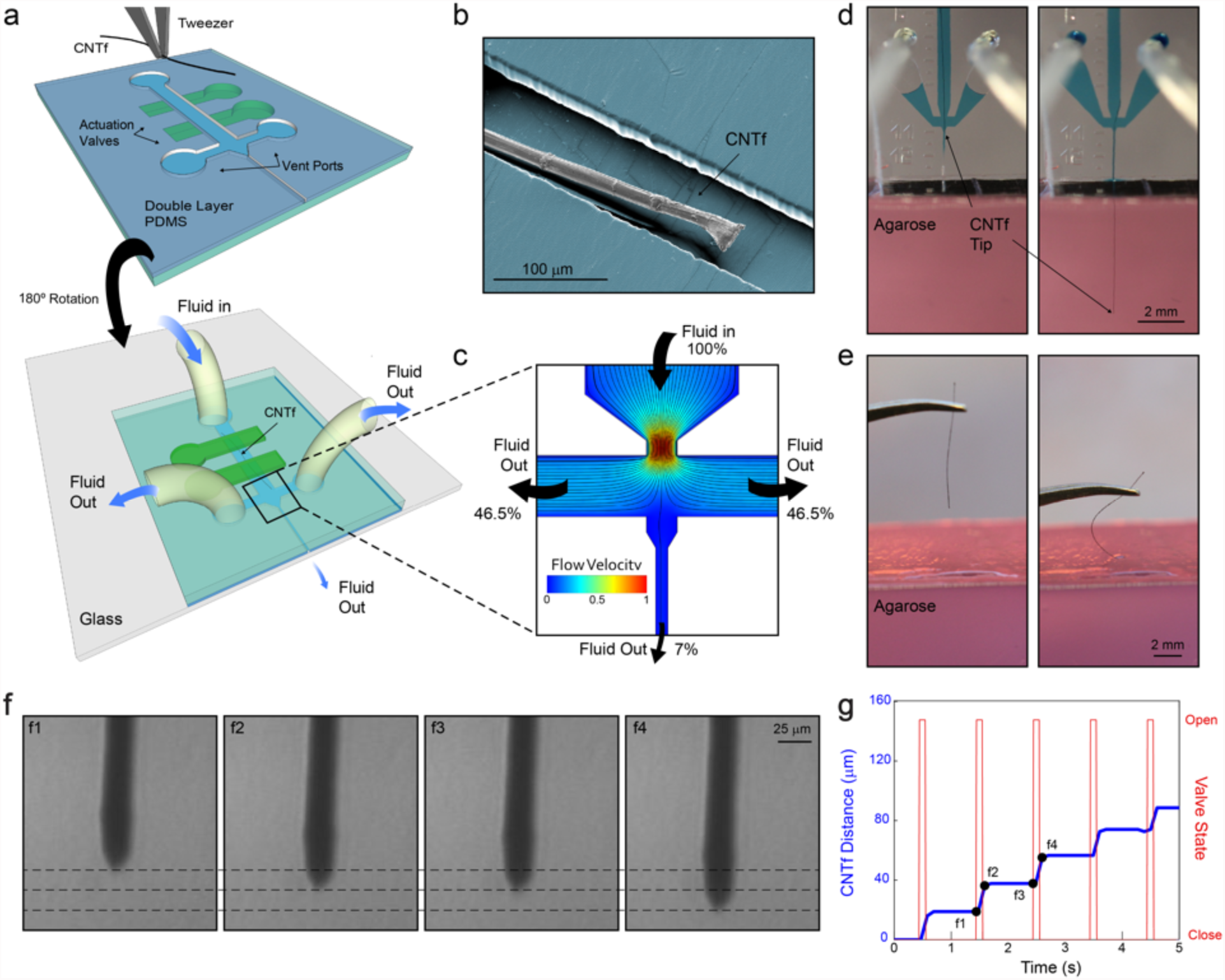
Device layout and microfluidic-assisted insertion of flexible CNTf microelectrodes *in vitro*. (a) Schematic of the two-layer PDMS microfluidic device. Microelectrodes are placed and aligned manually inside the channel (top). The device is then inverted and bonded to a glass substrate (bottom). Push-down actuation valves (green) provide on-chip flow control. (b) False-colored SEM image of a 12 μm diameter microelectrode inside the PDMS channel. (c) Velocity field and flow pathlines in the microfluidic device. More than 93% of the total volume of fluid injected is deviated to the side venting ports, which minimizes the amount of fluid delivered to the outlet channel. (d) Microfluidic-assisted insertion of 12 μm microelectrode in a brain phantom: the drag force produced by the fluid drives the fiber 4.5 mm into the phantom without evidence of bending. When mechanical insertion is attempted (e), the microelectrode irreversibly buckles upon contact with the agarose surface and does not penetrate inside the phantom. (f) Microscope images of the microelectrode position in an agar tissue phantom at times corresponding to panel g. Dashed lines, spaced approximately 15 μm, indicate the position of the fiber end. (g) Stepwise electrode insertion (blue trace) is controlled by opening the flow control valve for 100 ms intervals (red trace). The average fiber displacement during the open valve period is 16.4 ± 6.4 μm (mean ± s.d., n = 43 steps, 2 trials). Positions labeled f1 to f4 refer to the images shown in panel f.

The liquid flowing in the microfluidic channel exerts a viscous drag force on the microelectrode due to the velocity differential that distributes the force applied to the fiber and holds it under tension. Finite element simulations show that the force at which the CNTf electrode buckles (the critical buckling load) increases by approximately 3 times when the load is distributed along the probe rather than applied to a fixed point near rear of the electrode. Thus, compared to traditional insertion strategies that concentrate the load on a single point, we expect that our microfluidic devices will allow us to apply more force to the electrodes without causing buckling.

We first tested our microdrive using an agarose brain phantom. The combined effect of the tension on the fiber and the confinement in the microfluidic channel allowed us to drive a parylene-coated 12 μm diameter CNTf more than 4.5 mm deep into a brain phantom, (Figure 1 d, Supplementary Video M1). In contrast, all of our attempts (n = 4 fibers) to implant 12 μm diameter CNTfs without the fluidic microdrive resulted in buckling upon contact with the phantom surface (Figure 1 e, SI Video M2). This buckling was observed despite using manually actuated translation stages to guide straight and vertical entry into the phantom.

### Drive Fluid Diversion

A major advantage of our fluidic microdrives compared to traditional syringe injection is the fact that we can minimize the volume of fluid injected into the brain and thereby reduce any risk of overpressure that could produce trauma to the brain tissue. By creating a large hydraulic resistance through the exit channel, we can divert the fluid flow into microfluidic vent channels that safely transport the fluid away from the injection site. Because the hydraulic resistance depends inversely as the third power of the channel cross sectional area,^36,37^ we created three main sections of our device where we adjusted the channel width to control the fluidic resistances: a wide, low-resistance upstream channel connected to the flow input port converging into a high-resistance outlet channel for microelectrode delivery and two low-resistance side ports. Details on the two channel geometries used in this work are reported in Supplementary Materials (Supplementary Figure S10).

Computational analysis of the flow field in the device (Figure 1 b) shows that the area of maximum flow velocity is concentrated in the converging nozzle and that the venting ports divert more than 93% of the input volume with the remainder of the fluid flowing downstream to the exit channel. Note that these relative fluid flow rates are computed in the absence of the electrode. With an electrode blocking much of the exit channel we expect the fluidic resistance to be higher and even less fluid to be ejected with the electrode.

### Precise Positioning

The hydraulic design and the control of the on-chip valves enable actuation and positioning of the CNTf. In the microfluidic channel, we can control the fiber velocity and direction by controlling the fluid flow rate. As expected, larger flow rates corresponded to more rapid microelectrode motion (Supplementary Figure S5 and Video M3). Targeting of specific brain regions often requires insertion of electrodes to a specific depth. Moreover, recording individual neurons *in vivo* often requires even more precise control to place microelectrodes into the proximity of neuronal cell body. Thus, we explored the ability to microactuate the electrode using on-chip actuation valves. In an agar tissue phantom, we observed that opening the microfluidic actuation valves at 100 ms intervals allowed us to reliably advance a CNTf with 25 μm in diameter with a step size of 16.4 ± 6.4 μm (n = 43 steps, 2 trials, Figure 1 f and g).

### Recording Neural Activity in *Hydra*

As an example of the fluidic microdrive implementation, we performed electrophysiological measurements in the freshwater cnidarian *Hydra (H. littoralis)*. *Hydra* are a compelling model organism because they are easily cultured in the laboratory and are known to generate compound action potentials that correspond to body contractions (contraction bursts, CBs),^38^ but their soft, deformable body makes them exceedingly challenging for conventional electrophysiology. To interrogate *Hydra,* we modified our microfluidic system to control the position of the animal body. In our experimental setup depicted in Figure 2 a, the working Ag/AgCl electrode is connected to the microfluidic device and makes electrical contact to a microelectrode with 12 μm in diameter through the conductive drive solution (dextran in PBS, 40% w/w) filling the channels, yielding a final impedance of 663.7 ± 127.0 kΩ (mean ± s.d., n = 3 devices). Throughout the experiment, we held the *Hydra* approximately 3 mm downstream from the microdrive exit channel by applying moderate negative pressure to trap channels located in a separate PDMS block (Figure 2 b). The pressure applied by these channels was sufficient to prevent the animal from moving along the chamber, but allowed the *Hydra* to contract, elongate, and nod. It is worth noting that despite the low levels of negative pressure used for trapping, we observed that some of the *Hydra* tissue was dissociated and pulled through the trapping channels. While this damage did not prevent stereotypical contractions and elongations during roughly one hour of immobilization, improved immobilization chambers may be necessary for longer experiments.

**Figure 2.**
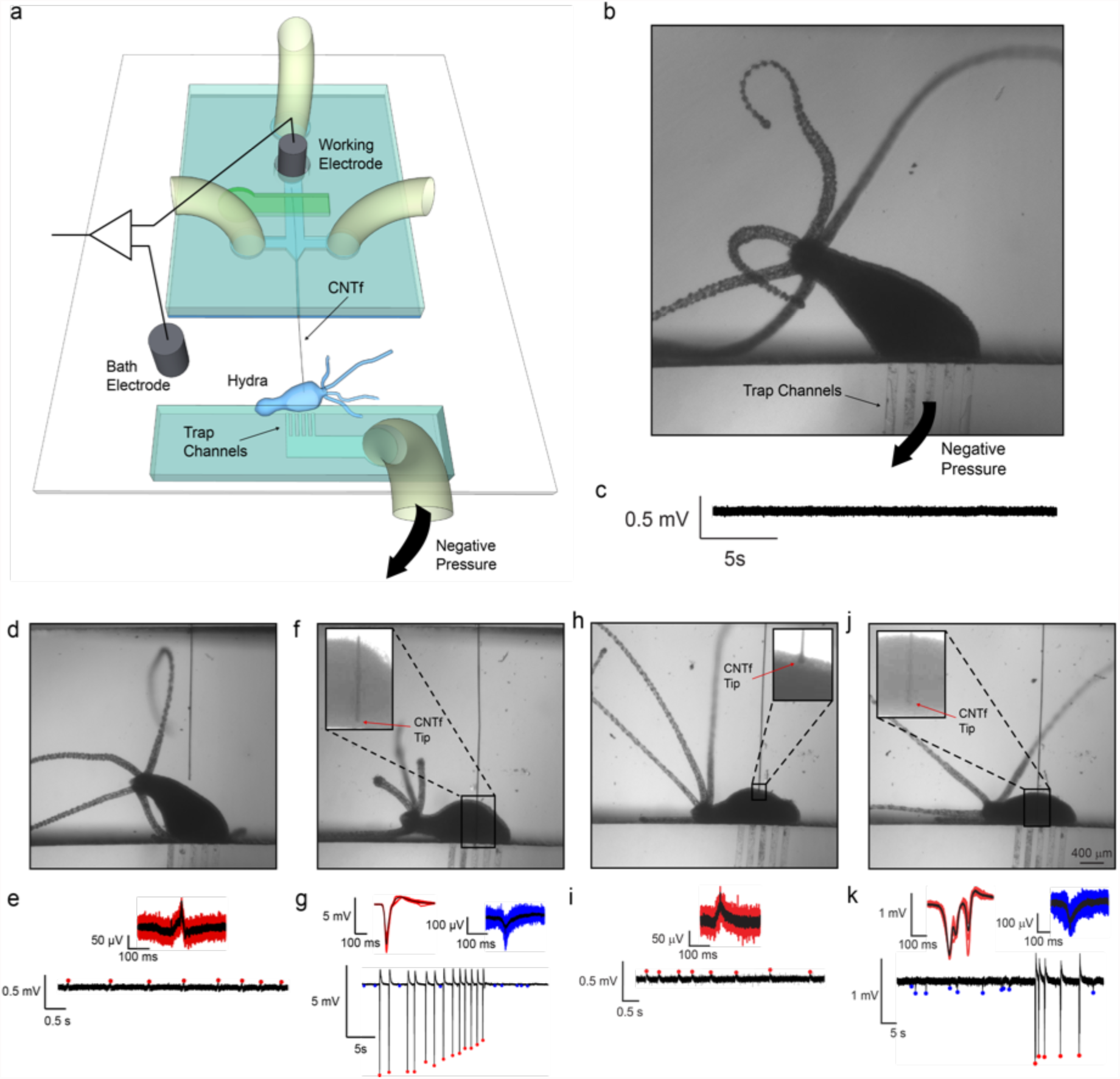
*In vivo* electrophysiology in *Hydra* (n = 3 animals). (a) Schematics of *Hydra* recording chamber. The reference electrode was placed in the bath while the working electrode was inserted into the microfluidic channel, making electrical contact with the CNTf microelectrode through the conductive Dextran solution. *Hydra* was secured ~3mm away from the microelectrode exit. (b) Optical microscope image shows *Hydra* trapped in the device by gentle negative pressure applied to the trap channels. (c) No peaks were observed when the microelectrode was more than 1 mm away from *Hydra* even during body contractions. (d) and (e) microelectrode located next to *Hydra* recorded small amplitude peaks (red) during body contractions only. (f) and (g) microelectrode inserted in *Hydra* recorded high amplitude peaks (red) during body contraction and small amplitude peaks (blue) in the absence of body contractions. (h) and (i) microelectrode was retracted to a position near the *Hydra* body. Small peaks were recorded only during body contractions similar to (e). (j) and (k) the microelectrode was re-inserted in the animal where we could once again record large peaks during body contractions (red) and small peaks in the absence of body contractions (blue) similar to panel (g).

By controlling the fluid flow and the on-chip valve actuation system, we were able to bring the microelectrode into contact with *Hydra* (n = 3 animals) and record compound action potentials. When the CNTf microelectrode was retracted into the fluidic device we recorded no electrophysiological activity even during body contractions (Figure 2 c); however, when we used the microfluidic device to position the microelectrode approximately 50 μm from the *Hydra* body, we detected small-amplitude spikes (~120 μV) that corresponded to body contractions (Figure 2 d, e and Supplementary Video M4). No activity was observed during body elongation and nodding. Next, we inserted the microelectrode into the *Hydra* and recorded spikes with much larger amplitudes (4.5 – 6 mV, Figure 2 f, g) correlated with body contractions. These large-amplitude spikes (a.k.a. contraction bursts, CB) are believed to originate from a nerve ring located at the junction of the tentacles and the body column. CBs correspond to synchronous neural and muscular activity, and the electrical recordings are likely a summation of this activity^39^. These spikes are consistent with previously reported CBs^40^ confirming that our fluidic-actuated CNTf microelectrodes act as microactuated flexible electrodes for electrophysiological recordings. In addition to large amplitude spikes, we also recorded low-amplitude spikes (50 –150 μV, Figure 2 g) that did not correlate with body contractions. These low-amplitude potentials are likely due to other behaviors such as elongation or tentacle contractions, which are known to correspond to small-amplitude spikes.^40^ It is worth noting that in previous studies, CBs were recorded with amplitudes of ~200 mV. These recordings were obtained using suction electrodes that seal the animal against a glass pipette,^40^ which greatly reduces the leakage current and increases signal amplitude. Recordings from implanted CNTf microelectrodes in *Hydra* are more similar to extracellular recordings in the brain, where no seal is formed, resulting in CBs with lower amplitude.

To illustrate our ability to precisely position the CNTf microelectrodes to record from different areas we used our fluidic microdrive to retract and re-insert the microelectrode. When we retracted the microelectrode, we once again recorded low-amplitude peaks correlated with contractions (Figure 2 h and i). After re-insertion, we recorded high-amplitude signals correlated with contractions together with the low-amplitude activity (Figure 2 k) independent from contraction. We noted, however, a change in the waveform of the contraction peaks. This variation in the waveform is likely due to a difference in the *Hydra* position, which agrees with previous reports that recorded CB waveforms depend on the position of the electrode with respect to the animal body.^40^

### Recording Neural Activity in Acute Brain Slices

Electrophysiology *ex vivo* in acute sections taken from the mammalian brain using manually-positioned glass pipette microelectrodes is a well-established experimental technique. To demonstrate that our fluidic microdrives can also place ultra-flexible electrodes at specific locations within a mammalian brain, we recorded neural activity in thalamocortical brain slices of mice (13-21 days old). The *ex vivo* slice provides the distinct advantage of allowing for the electrode position to be precisely visualized during the experiment, enabling assessment of the placement accuracy of our fluidic actuation technology. To minimize damage to the neural tissue, we etched the ends of 25 μm in diameter CNTf microelectrodes to create a sharp tip (30 degrees) using a FIB mill (Supplemental Figure S6). Using our fluidic microdrives we successfully inserted microelectrodes into cortex (up to ~1 mm deep, Figure 3 a, d) and into specific thalamic nuclei (~4mm from the cortical surface, Figure 3 a, e).

**Figure 3.**
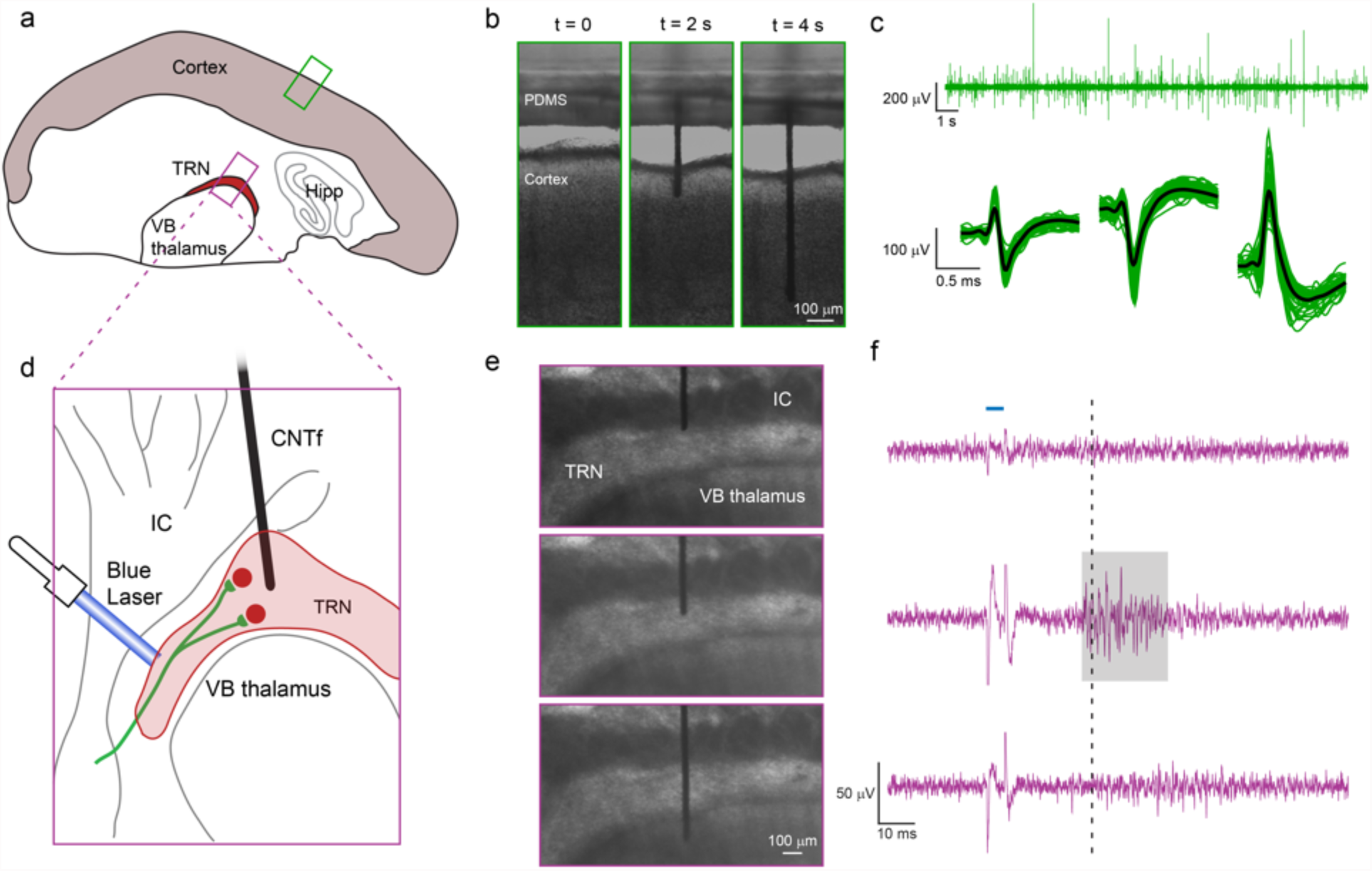
Microfluidic actuation and interrogation of CNS neural circuits in brain slices. (a) Schematic of thalamocortical section of a mouse brain depicting the two regions interrogated: Cortex and thalamic reticular nucleus (TRN). (b) Sequence of CNTf microelectrode insertion into neocortex. (c) Cortical activity recorded by the microelectrode in cortex (n = 3 slices, one slice per animal) (top). Automatic detection algorithms isolated action potentials from individual neurons, shown temporally aligned and averaged (bottom). (d) Schematic showing ChR2 expressing cholinergic synaptic afferents in ChAT–ChR2–EYFP mouse line (green) targeting TRN, but not the adjacent ventrobasal (VB) thalamus or internal capsule (IC). (e) and (f) Stimulation of cholinergic afferents with (5 ms) single laser pulses (blue horizontal line, F, top) triggers action potential activity specifically in TRN neurons. Spikes in the voltage recording during the optical stimulus (f) is an artifact produced by illuminating the microelectrode. When the microelectrode was positioned immediately above the TRN (e, top) no laser-evoked activity was recorded (f, top). Once the microelectrode was positioned within the TRN (e, middle), we recorded laser-evoked responses (highlighted by gray area) approximately 30 ms (dashed line) after stimulation. By inserting the microelectrode further, we could position the electrode in the VB thalamus (e, bottom), where no laser-evoked activity was detected (f, bottom), indicating that neuronal activity was acquired only from regions close to the tip of the microelectrode. Recordings in Cortex (b and c) and TRN (e and f) were obtained from slices from wild-type and transgenic mice, respectively.

Following microelectrode insertion into cortex (Figure 3 b) of brain slices extracted from wild type mice (n = 3 slices, one slice per animal), we were able to detect spontaneous neuronal activity as extracellular action potentials (Figure 3 c, top). Our automated spike detection and clustering algorithm isolated spikes with amplitudes around 50 μV to 500 μV (Figure 3 c, bottom).

Taking advantage of the possibility to precisely position the microelectrode with the fluidic microdrive, we moved the microelectrodes to record spatially confined neuronal activity in the thalamic reticular nucleus (TRN) of brain slices from transgenic mice. The TRN is a shell-like structure that in our slice preparation had a thickness of 168 ± 62 μm (n = 4 slices). Neurons in the TRN are the target of cholinergic synaptic afferents from the basal forebrain and the brainstem. Previous studies have shown that stimulation of these afferents leads to the fast and reliable activation of both nicotinic and muscarinic acetylcholine receptors (nAChRs and mAChRs)^41^ and the generation of short-latency action potentials specifically in TRN neurons, but not in neighboring thalamic nuclei.^41^ To selectively activate cholinergic afferents using optogenetics, we used brain slices obtained from ChAT–ChR2–EYFP transgenic mice (Figure 3 d), which express channelrhodopsin-2 (ChR2) specifically in cholinergic neurons.

When we positioned the microelectrode at the TRN/internal capsule (IC) boundary (Figure 3 e, top), we did not detect neuronal activity in response to optical stimulation (Figure 3 f, top). By opening the microfluidic flow control valves for 100 ms intervals, we then gently pushed the microelectrode into the TRN (Figure 3 e, middle). In this region, we observed action potential activity approximately 30 ms following laser stimulation (Figure 3 f, middle). Since neurons in the TRN do not express ChR2, the activity recorded in this region is the result of a postsynaptic response, and ~30 ms latency is in accordance with the literature.^42^ Finally, we pushed the microelectrode deeper into the ventrobasal (VB) thalamus below the TRN (Figure 3 e, bottom). In this configuration, no responses were recorded by the electrode (Figure 3 f, bottom), indicating that electrical activity was only detected by the microelectrode tip. To confirm that laser-evoked activity in the TRN was evoked by the activation of postsynaptic nAChRs, we bath-applied the specific nAChR antagonist dihydro-β-erythroidine hydrobromide (DHβE), which completely eliminated TRN activity (Supplementary Figure S8). Taken together, these experiments show that our platform can not only insert bare flexible electrodes into the cortex, but can also drive these electrodes deep within the mammalian brain and accurately position them in specific brain region.

### Recording Neural Activity *in vivo*

Next, we sought to demonstrate that the fluidic microdrive can be applied to *in vivo* rodent experiments. Using the microfluidic device, we successfully implanted flexible (12 μm diameter, n = 3 animal and 22 μm, n = 3 animals) CNTf microelectrodes to a maximum depth of approximately 4 mm (12 μm diameter: 3.75, 3.39, and 4.17 mm; 22 μm diameter: 4.13, 4.27, and 4.10 mm) into the brain of anesthetized rats (Figure 4 a and b, and Supplementary Movie M5). Currently, the maximum insertion depth is limited by the total length of the microfluidic channel, which is 10 mm for the devices we used in vivo. By using longer microelectrodes and longer flow channels, the drag forces can be increased and we expect to be able to drive CNTfs to even deeper regions *in vivo*.

**Figure 4.**
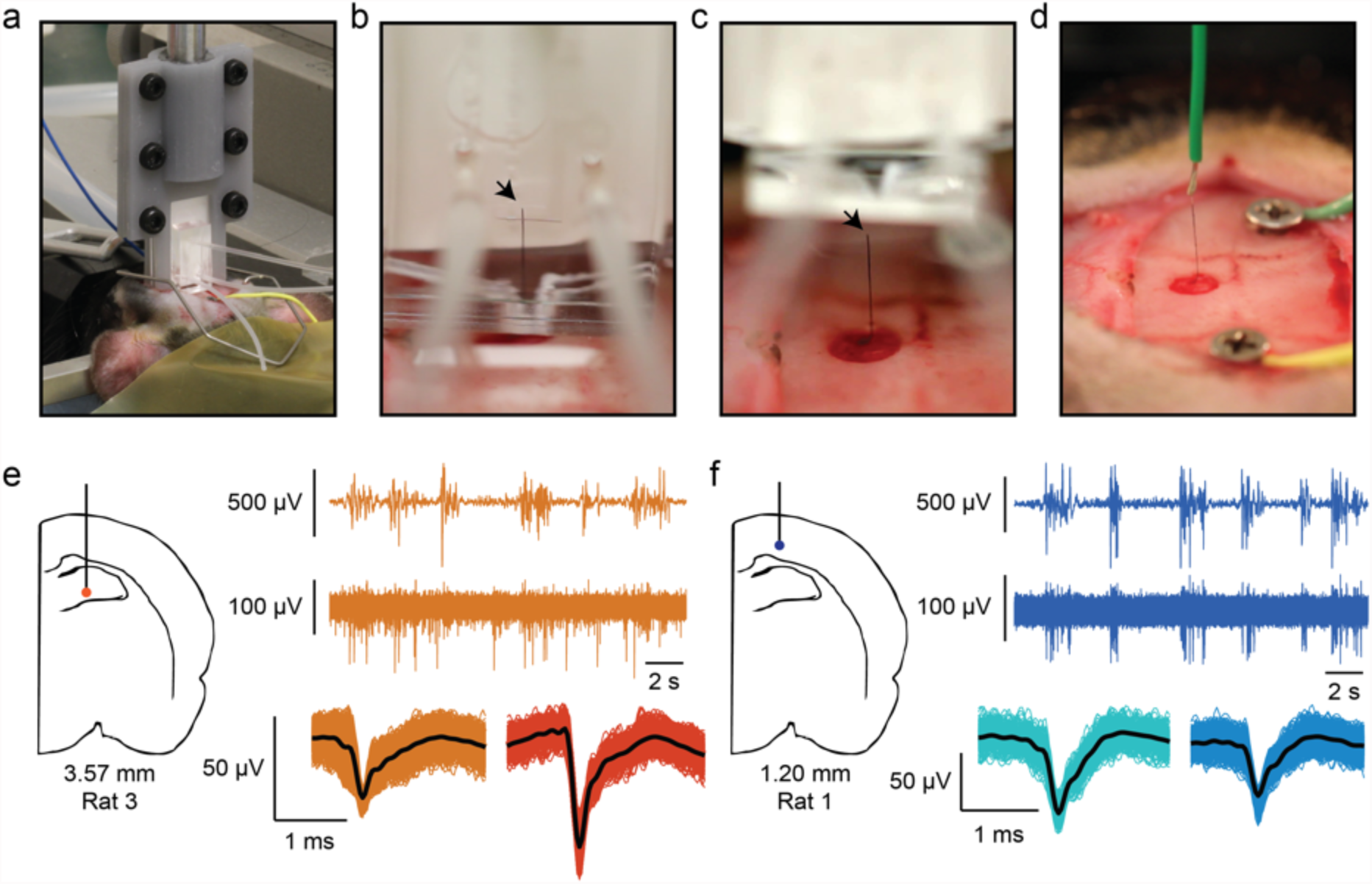
CNTf microelectrode recordings in anesthetized rats. (a) Photograph shows our microdrive attached to the stereotaxic arm using a 3D printed holder and positioned on top of the animal head. The device is gently placed in contact with the cortical surface through the craniotomy by actuating the stereotaxic arm. (b) Photograph of microelectrode following fluidic insertion into a rat brain. (c) To make electrical contact, the fluidic drive is retracted and (d) silver paint used to connect the microelectrode to a wire. Arrowheads mark the back end of the microelectrode. Note that (b-d) show a 22 μm diameter microelectrode for ease of visualization. (e and f) A representative example of 12 μm diameter microelectrode recordings at two depths from two different rats. (e) Orange traces are collected from 3.57 mm ventral, and (f) blue traces are from 1.20 mm ventral (as measured from the cortical surface). In each color, the top trace represents EEG signal collected from a screw placed over right frontal cortex. The burst-suppression pattern visible in the EEG is typical of isoflurane anesthesia. The bottom trace shows spikes recorded from the microelectrode, and clustered spikes are shown below that. Note that at 3.57 mm from cortex, spikes are not well timed to EEG bursts, whereas at a depth of 1.20 mm, spikes are tightly linked to EEG bursts.

As our goal is electrode insertion while minimizing fluid injection, we quantified the expected fluid released during insertion based on the total input volume. We measured the input drive fluid to be 14 ± 6 μL (n = 3 animals, 22-μm-diameter CNTf), which corresponds to an output volume of only 0.50 ± 0.23 μL, using the 3.5% ejection fraction estimated from our experimental data (see Supplementary Figure S9 and Methods). Thus, the output volume of our microdrive is 3 orders of magnitude smaller than a previously reported safe threshold for rats (up to 100 μL of solution^5^), greatly minimizing any potential damage associated with an increase in intracranial pressure. Also, it is worth noting that fluid leaving the microdrive is not injected into the brain, but instead forms a droplet surrounding the exit port (see Supplementary Figure S9). In a surgical setting, most of this dextran is quickly washed away by the ambient saline required to keep the brain wet when it is exposed, and any direct dextran contact with the brain is likely quickly diluted and washed away.

To record from the electrodes implanted into anesthetized rats, we retracted the microdrive (Figure 4 c) away from the brain following fluidic implantation and used Ag paint (Silver Print II, GC Electronics) to connect the exposed end of the CNTf microelectrode to a recording wire secured to a stereotaxic arm (Figure 4 d). By slowly moving the stereotaxic arm up from the brain we retracted the implanted microelectrode and recorded *in vivo* brain activity from a variety of depths (Figure 4 e and f). We implanted two skull screws, one as an electrical reference and a second as a monitor of cortical EEG activity. Under deep isoflurane anesthesia, cortical EEG cycles between periods of isotonic signal known as ‘suppressions’ and spindles of activity known as ‘bursts’.^17^ In burst-suppression, cortical firing is tightly linked to bursts, while other brain regions respond differently.^43-46^ Therefore, the temporal correlation between spike activity and bursts can indicate if the recorded neural activity is from cortical or sub-cortical regions.

In our recordings, we noted that spiking activity varied with the depth of our probe as measured from the cortical surface. At depths that suggest a sub-cortical, hippocampal recording, spiking activity is not well linked to bursts detected in the EEG (Figure 4 e), whereas at depths consistent with cortex, we observed spike timing tightly linked to bursts (Figure 4 f). Figure 4 shows data from 12 μm diameter microelectrodes, and recordings looked similar between 12 and 22 μm diameter microelectrodes. To verify that the implanted probe traveled straight down and did not deviate after entering the brain, we implanted a 22 μm microelectrode into an additional rat (n = 1 animal) using the fluidic microdrive. We then perfused the rat and sliced the brain on a cryostat to view the length of the microelectrode within the brain. We observed that the microelectrode traveled a straight trajectory through hippocampus (Supplementary Figure S11).

## Discussion

The fluidic microdrives introduced here enable implantation and actuation of flexible cellular-scale electrodes without using stiffeners or shuttles that would otherwise increase acute damage caused by the larger electrode footprint during implantation. In addition, our devices open up new possibilities for flexible electrodes by allowing precise electrode positioning to target locations within model organisms and the mammalian central nervous system. Compared to syringe injection, our fluidic microdrives nearly eliminate the volume of fluid injected into the brain, thus reducing potential damage from increased intracranial pressure. In addition, our approach requires no needle or delivery vehicle to penetrate the brain, potentially reducing damage to the tissue and blood brain barrier. While this reduced electrode footprint enabled by fluidic microdrives is expected to produce less acute damage, future work is needed to compare the long-term benefits for chronic *in vivo* recordings compared to conventional implantation approaches. Because our fluidic microdrives are compatible with conventional microfluidic devices, our technology could be integrated with high-throughput microfluidic chips for studying small model organisms like *Hydra* or *C. elegans*,^47-50^ providing an on-chip technology to precisely position stimulation and/or recording electrodes into specific regions of the animal.

We envision that fluidic microdrives can be employed for a variety of flexible neural probes and may become the preferred delivery technology. This study focused on CNTfs as model flexible electrode; however, the PDMS replica molding technique allows us to easily optimize the device design for a variety of probe sizes, materials and geometries. In order to scale up from one recording channel, flexible multi-channel probes with footprint similar to CNTfs can be fabricated using Parylene, Polyimide, or SU-8.^2,27,32^ Because the fluid ejected by the device is 3 orders of magnitude below the reported limit for fluid injected into the brain^5^, and this fluid is likely to be simply dispersed over the brain surface, we expect to be able to insert hundreds of electrodes without any adverse effects of the dextran drive solution. We also envision increasing the number of implanted electrodes by creating microdrives with multiple fluidic channels that can deliver numerous independent multi-channel electrodes. By eliminating the need for shuttles and mechanical methods for microactuation, it may also be possible to bring adjacent probes much closer together for high-density neural interfaces. The combination of emerging wireless, flexible, multi-electrode probes^4,5^ with high-density fluidic microdrives opens up exciting opportunities for chronic interfaces to large neuronal populations.

## Materials and methods

### Fabrication of CNTf microelectrodes

CNT fibers were fabricated from Meijo EC1.5-P CNTs (Meijo Nano Carbon Co., Ltd.) using a wet-spinning process previously reported.^14^ Recordings from *Hydra* were obtained using CNTf coated with a dielectric double layer of Al_2_O_3_ and HfO_2_ by Atomic Layer Deposition (ALD). To obtain a conformal coating, 12 μm diameter CNTfs were suspended between two glass slides separated by a few millimeters to prevent most of the fiber length from being in contact with a substrate. First, the fibers were exposed to an oxygen plasma cleaning process for a few minutes to clean residues and facilitate dielectric nucleation. Next, a standard ALD deposition (Cambridge Ultratech Savvanah S200) was performed. Al_2_O_3_ layer (50 nm) was grown using the precursors trimethylaluminum (TMA) and water and HfO_2_ layer (25 nm) used tetrakis[dimethylamido]hafnium (TDMAH) and water. The process chamber was at 150°C and 1 Torr pressure with N_2_ flowing at 20 sccm as carrier gas. Recordings from slices *ex vivo* and rat brain *in vivo* were obtained using Parylene coated fibers. Parylene C coating was preformed following standard protocols on a SCS Labcoter^®^ 2 Parylene Deposition System. A donut shape acrylic holder was specially designed to load the fiber in the coating chamber allowing most of their length to be suspended. Scanning electron microscope (SEM) inspection indicates conformal and crack-free coating (Supplementary Figure S1).

### Bending stiffness measurements

Bending modulus of insulated CNTf microelectrodes was measured using cantilever experiments,^51^ using a similar setup to what described by Clapp et al.^52^ CNTf samples of approximately 25 cm in length were hung vertically with a mass ~ 1.3 g in an oven at 115 °C for at least 4 hours. The fibers were then allowed to cool under tension for 6 hours in a glove box. This annealing process under tension was used to straighten the fibers and eliminate any bending history in them. Later, fibers were coated with parylene while still under tension. The straight insulated fibers were placed on a Teflon block and both ends were cut using sharp razor blade. One end of the fiber was clamped and opposite end was allowed to hang under self-weight as a cantilever in a glove box to minimize errors from air current disturbances. The fibers were slowly pulled backwards decreasing the cantilever length until no deformation was observed. Photos of the cantilever were taken at different lengths using a Nikon Coolpix 900 digital single-lens reflex camera mounted on a tripod. The cantilever profiles were then mathematically processed using Matlab software and bending stiffness was calculated from fitting the measured deformation according to the *elastica* model, accounting for geometric nonlinearities. Results of bending stiffness measurements on CNTf microelectrodes and comparison with other neural probes are reported in Supplementary Figure S3. Remarkably, due to relative shear sliding of the CNTs forming the fiber, the bending stiffness of CNTfs is not directly associated with the moment of inertia of the cross sectional area, even if it is here referred to as *EI* following the classical notation.^51^

### Leakage current tests

The insulation coating thickness was measured with an ellipsometer (Filmetrics). The coating morphology was observed using both optical and electron microscopy to check for any defects. Furthermore, integrity of insulation coatings was assessed by measuring DC leakage currents and electrochemical impedance spectroscopy (EIS). Leakage currents were measured between insulated CNTfs (1 cm length) and a large carbon wire serving as counter and reference electrode at room temperature^53^ on 1 cm of CNT fiber in PBS. A Gamry Reference 600 potentiostat (Gamry Instruments, input impedance 100 TΩ) was used to apply DC voltage bias of 1 V and the leakage current between the insulated CNT fibers and the counter electrode was measured. EIS measurements were performed with 20 mVpp input voltage in the range 1 Hz-100kHz between coated CNTf and the carbon counter electrode. Ag/AgCl electrode served as reference. To evaluate the mechanical stability of the coatings, we also measured leakage currents after the CNTf samples were bent at 5 different points, spaced 1 to 2 mm apart. CNTf were bent at 90^o^ by pressing them hard at a point and bending it across the point using tweezers. The bent fibers were immersed back in PBS and the leakage current measurement protocol was repeated. Negligible difference in the leakage current values before and after the bending of insulated CNT fibers suggest the mechanical stability of parylene coatings.

### FIB patterning CNTf microelectrodes

To facilitate insertion and reduce damage in tissue, parylene coated CNTf with 22 and 25 μm in diameter were sharpened using standard FIB (FEI Helios NanoLab 660 DualBeam) protocols. A 30kV gallium beam (65nA) was focused on the surface of the fiber to cut it at a 30° angle with respect to its axis.

### Dextran rheology

Dextran viscosity measurements were carried out at room temperature with Advanced Rheometric Expansion System (ARES, Rheometrics Scientific, now TA Instrument) in a Couette geometry. A solution of 40 % w/w Dextran in DI was loaded in a titanium cup (diameter = 16.5 mm), which rotates relative to a titanium bob in the center of the cup. Viscosity was measured as a function of shear rate (test duration: 20 min, shear rate: 0.1 to 100 s^-1^).

### Microfluidic device fabrication

Double layer PDMS microfluidic devices were fabricated following standard soft-lithography technique. Briefly, Si wafers (University Wafer) were spin coated with SU-8 2050 (Microchem), followed by a photolithography process which defined valve and flow channels on two separate wafers. Next, a thin layer (approximately 70 μm) of flexible (20:1 Elastomer:Cross-linker, w/w ratio) PDMS (RTV615, Fischer Scientific Company LLC) was spin-coated on the flow channel patterned wafer. At the same time, a thick (approximately 2 mm, 5:1 Elastomer:Cross-linker, w/w ratio) PDMS was poured on the valve layer patterned wafer. Both wafers were baked in an oven at 90° C: 15 min for the flow layer and 45 min for the valve layer. After alignment of the flow and valve control layers, the devices were baked for additional 24 hours at 90°C.

CNTf microelectrodes with diameters of 12 μm (agarose, *Hydra*, and *in vivo* experiments), 22 μm (*in vivo* experiments) and 25 μm (CNTf actuation and slices experiments) and length between 7 and 10 mm were manually loaded in the PDMS device using carbon-tip tweezers. To seal the microfluidic device with the fiber, an oxygen plasma cleaning step (Harrick plasma PDC-001) on the PDMS and glass was performed, allowing for a strong and leak free covalent bound between their exposed surfaces. Each microelectrode was used only once per trial and never reused.

We found that overall device yield was approximately 80% with the most common failure mode being PDMS delamination. Plasma binding between PDMS and glass is extensively used in microfluidic devices operated under low pressures; however, the high input pressure necessary to produce the high flow rates that drive microelectrodes in tissue can cause PDMS delamination. This delamination can produce undesirable leaks that reduce the velocity of the fluid flow. We expect that the inconsistency in the plasma binding may be due to the cleanliness of the PDMS surfaces or variability in the plasma density and gas ratios, and that improving these elements may increase device yield.

### Measurement of ejected dextran volume

The volume of fluid ejected from the device through the exit channel was obtained by measuring the size of the dextran droplet formed at the output channel in the device geometry used for *in vivo* experiments. With no microelectrodes in the channel, we flowed 4.1 ± 2.5 μL (n = 5 devices) of dextran solution through the device and observed a droplet form at the output channel. By modeling this droplet as a fraction of a sphere (see Supplementary Figure S9) we estimated the volume fluid to be 3.5 ± 1.5% (n = 5 devices) of the input volume, which is lower than the theoretical value of 7% estimated from CFD simulations. This difference is likely due to variations on the channel geometry during device fabrication relative to the ideal computational model.

### Brain phantom insertion tests

Agarose gel was used as brain phantom for insertion tests (Fig. 1 d and e). The gel was prepared by first mixing agarose (Sigma-Aldrich Corporation) with DI water on a 0.6% w/w concentration.^54^ The solution was then heated in a microwave until boiling and finally left at room temperature to gelate at least 2 hours. For imaging purpose, a drop of red food colorant was added to the solution.

Microfluidics devices were loaded with parylene-coated 12 μm diameter CNTf microelectrodes as described above. The glass slide was vertically mounted on a micromanipulator and gently brought in contact with the agarose gel. To insert the fiber in agarose, manual pressure was applied to the syringe filled with a blue Dextran solution (40%w/w in DI with blue food colorant) connected to the fiber input port (Supplementary Video M1).

Manual CNTf insertion attempted by moving a similar fiber vertically towards the agarose gel using a self-closing tweezer assembled on a micromanipulator. The CNTf was held approximately 4 mm away from the edge and the micromanipulator was manually operated to move the fiber at an average speed of approximately 0.3 mm/s (Supplementary Video M2).

### Tissue phantom actuation test

Microfluidic devices loaded with FIB-sharpened, 25 μm diameter CNT fibers were fabricated as described above and horizontally attached in a petri dish. Valve channels were filled with DI water and connected to a valve controller. A Matlab (Mathworks Inc.) script was used to switch between Closed and Open valve states. A cubic block of agar gel (0.6 % w/w in DI prepared as described above) approximately 1 cm^3^ was gently placed in contact with the exit channel of the device. To drive the fiber, manual pressure was applied to the syringe filled with Dextran solution (40% w/w in DI) connected to the fiber input port. A video of the fiber movement in agar was collected at approximately 8.9 fps using Hamamatsu Orca 03-G camera mounted on an inverted Nikon scope. A Matlab script was used to isolate the fiber from the background and determine the position of the distal end.

### Device impedance in Hydra and brain slice experiments

To measure impedance of the devices used in *Hydra* and brain slices experiments, the CNTf microelectrode was inserted in the fluidic microdrive and connected on one end to the working Ag/AgCl electrode through the conductive Dextran solution, which is made by mixing Dextran with PBS. The distal end of the microelectrode was advanced inside a 1x PBS bath. A Ag/AgCl immersed in the PBS bath was used as a reference and counter electrode.

### *Hydra* culture

*Hydra* (*H*. *littoralis* obtained from Carolina Biological Supply Company) were cultivated in plastic vials at room temperature in standard *Hydra* culture medium (protocol adapted from Steele lab). The animals were fed once every two days with freshly hatched brine shrimps (*Artemia naupli*). Before electrophysiology experiments *Hydras* were food deprived for 24 hours.

### *Hydra* electrophysiology

A glass slide containing both the fluidic microdrive and the *Hydra* trap was fixed to a plastic Petri dish filled with *Hydra* media. *Hydra* (n=3) with length between 1.5 and 2.0 mm were randomly selected and partially immobilized by the *Hydra* trap. This was accomplished by positioning the animal using a glass pipette next to the trap channels, followed by application of negative pressure at the trap port. Next, 12 μm in diameter CNTf microelectrodes (138 ± 33 kΩ, microelectrode impedance, n = 3 CNTfs, final device impedance 663 ± 127 kΩ, n = 3 devices, 10 mV, 1 kHz) were actuated towards the *Hydra* by applying manual pressure to a syringe connected to the fiber input port and filled with a conductive dextran solution (40% w/w in 1x PBS). Valves were actuated (opened and closed) to guarantee precise positioning of the fiber inside or next to the animal.

The working Ag/AgCl electrode was placed in contact with the conductive Dextran solution in the microfluidic channel while the reference Ag/AgCl electrode was placed in the *Hydra* medium. Data were collected by using an AM Systems model 1800 amplifier (input impedance 100 MΩ) and a Digidata 1550 digitizer (Molecular Devices) at 10 kHz sampling rate with 0.5 Hz low-cut filter and 20 kHz high-cut filter.

### Fluid dynamic analysis

Flow simulations were performed using Comsol (Comsol Inc.). Stationary Navier-Stokes equation was solved in laminar flow regime using a finite element method applied to a 3D model of the device layout. Boundary conditions were set to constant pressure of 10 atm at the inlet port and 1 atm at the outlet port and no-slip on the sidewalls.

### Buckling analysis

Critical buckling load of microelectrodes was obtained by solving an eigenvalue-based linear buckling analysis under stationary conditions using Comsol (Comsol Inc.). CNTfs were modelled as Euler-Bernoulli beams with 1 mm in length and 12 μm in diameter. A fixed constraint was set to one of the extremes and a unitary load directed towards this fixed boundary was applied. This load was either concentrated on the free extreme or distributed along the CNTf length. These calculations show that the critical buckling load for the distributed load scenario is approximately 3 times higher than the concentrated load case.

### Brain slice preparation

Young mice (male and female) were randomly selected from two lines (P13-21): C57BL/6J wild-type (n = 3 animals) and bacterial artificial chromosome (BAC)-transgenic mice (n = 3 animals) expressing channelrhodopsin-2 (ChR2) under the control of the choline acetyltransferase (ChAT) promoter (ChAT–ChR2–EYFP). Wild-type and transgenic animals were used for recordings in cortex (Figure 4B and C) and in TRN (Figure 4e and f), respectively. Thalamocortical slices (400 μm) were prepared as described previously.^55^ Animals were anesthesized with isofluorane and decapitated, following procedures in accordance with NIH guidelines and approved by the UTHealth animal welfare committee. Slices (450 μm) were cut in an ice-cold solution consisting of (in mM): 234 sucrose, 2.5 KCl, 1.25 NaH2PO4, 10 MgSO4, 26 NaHCO3, 10 glucose, and 0.5 CaCl2, saturated with 95% O2-5% CO2, using a vibratome (Leica VT1200S) at speeds of 0.2 mm/s and a blade vibration amplitude of 0.8 mm. Slices were incubated at 34°C for 40 min in artificial cerebrospinal fluid (ACSF) containing (in mM): 126 NaCl, 26 NaHCO3, 2.5 KCl, 1.25 NaH2PO4, 10 glucose, 2 CaCl2, and 2 MgCl2. Slices were then kept at room temperature prior to recordings.

### Brain slice electrophysiology

Fluidic microdrives were sealed at the edge of a 500 μm thick glass slide and mounted on a 60 mm-diameter plastic petri dish. Tubes for ACSF perfusion were placed on opposing sides of the dish. Freshly dissected 450 μm brain slices were placed on a 150 μm thick glass slide pre-coated with poly-L-lysine (Sigma Aldrich). The brain slice was manually positioned as close as possible to the device with cortex facing the fiber exit channel. All recordings were carried out at near-physiological temperatures (32-34 °C).

CNTf microelectrodes (25 μm diameter, 355 ± 93 kΩ microelectrode impedance, n = 3 CNTfs, final device impedance 261 ± 16 kΩ, n = 2 devices, 10 mV, 1 kHz) were inserted into brain slices by applying manual pressure to a syringe connected to the fiber input port and filled with a highly conductive Dextran solution (40% w/w in 20x PBS). Valves were actuated (opened for 100ms and closed for 900ms) to guarantee precise positioning and actuation of the fiber inside the brain slice. For cortical recordings, the microelectrode was actuated until spikes were visible in the electrical recording (n = 3 slices, each from different animals).

The working Ag/AgCl electrode was placed in contact with the conductive Dextran solution in the fiber channel while the reference Ag/AgCl electrode was placed in contact with the ACSF bath in the dish. Data were collected by using an AM Systems model 1800 amplifier and a Digidata 1550 digitizer (Molecular Devices) at a 50 kHz sampling rate with 0.5 Hz lowpass filter and 20 kHz highpass filter.

In all experiments on brain slices extracted from transgenic mice, when the microelectrode tip was in the TRN we recorded light-evoked activity (n = 3 animals). For microelectrodes with thin parylene insulation layers (1.2 μm), we also observed light-evoked activity after we actuated the tip through the TRN leaving the insulated portion within the TRN (n = 2 animals). To prevent electrical recording through the insulating layer we increased the parylene coating thickness to 3.3 μm. Under this condition, we recorded activity exclusively from the tip (Figure 3e and f, n = 1 animal). Thickness of TRN was measured in n = 4 brain slices used for experiments. The mean thickness was calculated by averaging the values of 3 regions in a given slice. Optical stimulation was performed using a DL447 laser (Crystalaser) at a wavelength of 440–447 nm. Single laser pulses of 5 ms duration and 5 mW power were used to activate cholinergic afferents in the TRN.

### Impedance to ground

To estimate the impedance to ground through the conductive Dextran solution (Z_Dex_, Supplementary Figure S7) during Hydra and brain slice recordings, we analyzed the impedance spectrum of devices loaded with microelectrodes (Supplementary Figure S12). We expect the device impedance to converge to Z_Dex_ at very low frequencies. At frequencies below 10 Hz we found that the impedance stabilized near 1.5 MΩ which we expect to be primarily the DC impedance through the dextran solution. This impedance to ground was roughly twice the interfacial impedance of the CNTf (Z_Dex_ ≈ 2Z_int_), which implies that the signal is attenuated by approximately 50%. Such attenuation, however, does not prevent us to record compound action potentials in *Hydra* and neuronal spikes in brain slices.

### Neural recording *in vivo* in rats

Long-Evans rats (1 male and 2 females between 4 and 21 months old for 12 μm diameter microelectrode, and 1 male and 2 females between 3 and 22 months old for 22 μm diameter microelectrode, randomly selected) were used for our acute *in vivo* recordings. All the procedures were approved and monitored by the Rice University Institutional Animal Care and Use Committee. Rats were given buprenorphine (0.02 mg/kg) then induced under 5% isoflurane and maintained at 1-2.5% isoflurane for the duration of surgery and recording. After the initial incision and cleaning of the skull, two screws were placed in burr holes made over right frontal cortex (roughly 1.5 mm anterior and 2 mm lateral to bregma) and over the cerebellar bone for use as an EEG screw and ground screw, respectively. Next, a craniotomy was drilled over the left parietal lobe centered at 2 mm lateral and 3 mm posterior to bregma and dura over part of the exposed brain was removed. The fluidic microdrive was attached to a stereotaxic arm, centered and lowered over the durotomy. Once lowered, fluid pressure was applied to the input port via a tube connected to a syringe filled with dextran. Once pressure at the input port began to flow the dextran solution through the microchannel, the microelectrode proceeded to move and was implanted into the brain over the course of 5-10 seconds (Supplementary Video M4). In practice, the fluid injection parameters are adjusted during the experiment based on the fiber displacement. Once the fiber begins moving we can adjust the flow rate to achieve the desired velocity and penetration depth by monitoring the fiber displacement in the microfluidic device using a microscope or magnification optics. After the implantation was complete, the microdrive was slowly retracted with the stereotaxic arm while slight positive pressure was applied to prevent the probe from being retracted with the device (Supplementary Video M6). Once the CNTf microelectrode was free from the microdrive, another stereotaxic arm with a recording wire attached to it was brought close to the exposed end of the microelectrode. Ag paint (Silver Print II, GC Electronics) was applied to the recording wire and then the recording wire and microelectrode were brought together bridged by the Ag paint. After allowing the Ag paint to dry for 5 minutes, *in vivo* impedance spectroscopy was performed in the two-electrode configuration, using the fibers as working electrode and the skull screw as a reference. Impedance spectra were acquired by applying a 10 mV AC voltage in the 100 mHz to 1 kHZ range with a Gamry 600+ potentiostat. The average impedance of the CNTf microelectrode used *in vivo* was 430 ± 140 kΩ (22 μm in diameter, n = 3 CNTfs) and 138 ± 33 kΩ (12 μm in diameter, n=3). Note that the 22 μm diameter CNTf was sharpened using FIB to reduce tissue damage. This process increases the electrode impedance likely due to local heating, which can lead to the formation of amorphous carbon that can increase the interfacial impedance.

The recording wire was retracted in steps of 150 to 500 μm and recordings were taken at each step. During retraction, the electrode was visually monitored under magnification, and absolute implantation depth for each recording was inferred from the depth at which the electrode exited the brain.

### Histological verification of *in vivo* implantation

One male Long Evans rat was implanted with a 22 μm diameter microelectrode using the fluidic microdrive as described above. Immediately after implant, the rat was transcardially perfused with isotonic sucrose followed by 4% paraformaldehyde (PFA) while under deep isoflurane anesthesia. The brain was then extracted from the skull without disturbing the microelectrode and left to post-fix in PFA for 24 hours, then cryoprotected in 30% sucrose until the brain sunk (about 48 hours). After cryoprotection, the brain was mounted on a cryostat (Leica Microsystems) and cut into coronal sections 60 μm thick until the microelectrode was visible.

### Brain slice and *in vivo* data analysis

Electrophysiological recordings were analyzed and processed off-line with a custom Matlab (Mathworks Inc.) script. To isolate spikes, the raw data stream was first digitally filtered between 300 Hz and 6 KHz. Spike detection was performed on the bandpass filtered data based on an amplitude threshold automatically set as 4.5 times the estimated standard deviation of the noise.^56^ Principal components of the waveforms were calculated on a 3 ms window isolated from the threshold-crossing spikes followed by unsupervised clustering and sorting.

## Acknowledgements

This work was supported by a DARPA Young Faculty Award to J. T. R. (D14AP00049), Welch Foundation (C-1668), the National Science Foundation (CBET-1351692), the Air Force Office of Scientific Research (FA9550-15-1-0370), the American Heart Association (15CSA24460004) and the National Institutes of Health (NS077989). The content of this report does not necessarily reflect the position or the policy of the Government, and no official endorsement should be inferred. The authors thank Colin Young for his help with CNTf spinning, and thank Robert E. Steele, Celina Juliano, Stephan Siebert, Christophe Dupre, and Rafael Yuste for useful discussions of *Hydra* physiology and care. F. V., D. G. V, M. P., C. K., and J. T. R. are authors of a provisional patent application describing the fluidic microdrive. J. T. R. and C. K. are co-founders of a company investigating commercial applications of fluidic microdrives.

